# Investigating age- and sex-specific effects of socialization on voluntary ethanol administration using a novel vapor paradigm

**DOI:** 10.1101/2020.11.02.364927

**Authors:** Christopher D. Walker, Hannah G. Sexton, Mary-Louise Risher

## Abstract

**Introduction:** Adolescence is characterized as a transitional developmental period between childhood and adulthood that is associated with increased freedom and novel experiences that are frequently peer-influenced. Due to newfound independence, there is a higher prevalence of alcohol consumption, which is heightened by the rewarding effects of alcohol. However, the contributions of social interaction and sexual dimorphism to alcohol intake are not fully understood. Here we explore the use a novel self-administration ethanol (EtOH) vapor system to investigate the sexual dimorphic nature of socially facilitated ethanol exposure.

**Methods:** Adolescent and adult male and female Sprague-Dawley rats underwent a novel voluntary intermittent EtOH vapor paradigm. Nosepoke initiated self-administration vapor chambers administered 20mg/L of vaporized EtOH or air into the chamber following each nosepoke. Beginning on postnatal day 30 (PND30), during the onset of adolescence, or 70 (PND70), at the onset of adulthood, animals were placed in vapor chambers for 4hr every other day for 40 sessions. All animals underwent 10 sessions with their cagemate (social access) followed by 10 sessions in isolation (isolated access), a 10-day forced abstinence period, 10 sessions isolated access, and 10 sessions social access.

**Results:** These data reveal that despite low EtOH consumption across all groups, adolescent (PND30) and adult (PND70) female rats voluntarily self-administered more EtOH vapor per body weight than age-matched males, while male rats increased EtOH preference over sessions regardless of age. In addition, all rats regardless of sex or age voluntarily self-administered more EtOH vapor per body weight during the social access session than during the subsequent isolated access sessions.

**Conclusion:** These data demonstrate that under these experimental parameters, male and female rats regardless of age do not self-administer high quantities of EtOH vapor using this paradigm. Further work is required to determine whether the nose-poke EtOH vapor self-administration apparatus can be modified to promote high voluntary EtOH consumption that can be socially facilitated. These data demonstrate that with further investigation, the self-administration EtOH vapor system could be an effective alternative to other methods of voluntary EtOH administration to further our understanding of socially facilitated drinking.

## INTRODUCTION

Excessive alcohol consumption is the leading cause of preventable death in the United States and contributes to health, social, and economic problems (Collaborators et al., 2018). It has been shown that lifetime prevalence of severe alcohol use disorder is 18.3% in men and 9.7% in women (Grant et al., 2015). With men showing significantly higher drinking frequency, quantity, and rate of binge consumption compared to women (Wilsnack et al., 2009). Despite men drinking more the women, excessive alcohol consumption in women is on the rise (White et al., 2015, Slade et al., 2016). A 2015 study showed that in 2010, excessive drinking accounted for one in ten deaths in the United States and cost more than $248 billion (Stahre et al.,; 2014, Sacks et al., 2015). It was also shown that underage alcohol consumption cost the U.S. $24.3 billion that same year (Sacks et al., 2015). A 2019 study revealed that 29% of high school students consumed alcohol and 14% did so in a binge-like fashion (Jones et al., 2020). When parsing out these statistics, they found that 31.9% and 26.4% of high school females and males (respectively) had consumed alcohol and 14.6% of these females and 12.7% of these males did so in a binge-like fashion (Jones et al., 2020).

Adolescence is the dynamic period of transition from childhood to adulthood that encompasses, but is not limited to, puberty. Adolescence spans the teenage years (Peterson et al., 1996) including the mid to late 20’s, which is often termed ‘late adolescence’ or ‘emerging adulthood’ (Arnett, 2004, Baumrind, 1987). It is during this time that individuals have less adult supervision allowing for more opportunities to experiment with drugs, alcohol, and sexual behaviors, and sex differences in alcohol consumption begin to emerge (Witt, 2007).

An important aspect of adolescent development is peer-peer interactions. Social interactions during this time play a vital developmental role as they are critical for shaping cognitive growth leading to a sense of ’self’ or the development of one’s own identity (Burnett and Blakemore, 2009, Sebastian et al., 2008). Through peer reinforcement, positive and/or negative, adolescents tend to adopt behaviors that reflect those of their social groups. For example, adolescents are more prone to participating in risky behavior, such as consuming alcohol, when interacting with their peers than when they are alone. This kind of peer reinforcement can be direct, via peer pressure or the offer to supply alcohol, or indirect, in which the adolescent models the behavior they perceive to fit the social norms of their chosen social group (Jackson et al., 2005, Borsari and Carey, 2003). Animal studies have revealed interesting results regarding how social dynamics that can influence alcohol consumption. Blanchard et al. (1987) demonstrated that subordinate rats within Long-Evans mixed sex colonies, consumed significantly more alcohol than dominant rats, possibly due to the anxiolytic effects of alcohol. Interestingly, the females within these colonies drank significantly more than the males. Similar findings have also been reported in Wistar rats (Wolffgramm and Heyne, 1991). However, these studies include the added component of anxiety and stress. How underlying factors that are not dependent on negative experiences contribute to changes in alcohol seeking in social situations still requires further investigation. Some work has been done in this area. A number of studies have shown that rats show increased preference for novel substances when they observe another rat being exposed to the same substance (Galef et al., 1997). One model in particular that has become increasingly valuable for studying the biological mechanisms of social interaction is the vole due to their unique ability to form life-long bonds and their willingness to consume alcohol at levels equivalent to C57BL/6 mice (Anacker and Ryabinin, 2010). Recent work has demonstrated that voles will adjust their alcohol consumption to match that of the animal that it is paired with (Anacker et al., 2011). However, modeling the human experience in which drinking is influenced when peers are present without the confound of stress or anxiety remains to be extensively explored in rats. Moving forward, it will be important to develop experimental models that allow researchers to quantify the impact of socializing during EtOH exposure, allowing researchers to better correlate laboratory observations with real world behavioral consequences.

Parallels are beginning to emerge between human findings and rat models of binge EtOH exposure, making them an appreciable model for conducting alcohol studies (Spear and Swartzwelder, 2014, Crews et al., 2019). Adolescent rats are similar to adolescent humans as they are less sensitive than adults to the motor impairing, aversive, and sedative effects of acute EtOH and more sensitive to the social facilitating and rewarding effects of EtOH (Anderson et al., 2010, Ramirez and Spear, 2010, Silveri and Spear, 1998, Varlinskaya and Spear, 2002). Interestingly, it has been shown that male adolescent EtOH exposure can result in the persistence of this adolescent phenotype in select behaviors. Adult male rats that have been exposed to EtOH during adolescents were found to be less sensitive to the motor impairing and sedative effects of EtOH (Silveri and Spear, 1998, Varlinskaya and Spear, 2002, Ramirez and Spear, 2010). Multiple labs have previously demonstrated that there are behavioral, cognitive, and electrophysiological changes that persist into adulthood after adolescent binge EtOH consumption in a male rat model (Crews et al., 2019, Spear and Swartzwelder, 2014). At the behavioral level, adolescent binge EtOH exposure has led to increased impulsivity (Gilpin et al., 2012), deficits in reversal learning (Coleman et al., 2014), and object recognition in adulthood (Montesinos et al., 2015). With regard to neuronal function, it has been demonstrated that adolescent binge EtOH exposure results in enhanced synaptic plasticity that was correlated to an increase in immature dendritic spines (Risher et al., 2015a). It was also shown that adolescent binge EtOH consumption results in loss of adult neurons in the hippocampal CA1 along with an increase expression of astrocyte specific synaptogenic factors and astrocyte reactivity (Risher et al., 2015b). In summary, this model has yielded substantial insight into the long-term consequences of binge-level alcohol consumption during adolescent development. However, there are limitations in the methodology due to the involuntary nature of EtOH consumption in rats.

Involuntary EtOH administration has been necessary because rats have an innate aversion to the taste of EtOH, making it difficult to replicate voluntary binge-level consumption that we see in humans. However, some studies have suggested that experimenter-administrated EtOH during adolescence may impact the motivation of the rat to voluntarily consume EtOH when assessing voluntary consumption in later studies. This has been demonstrated using involuntary exposure to EtOH via injection which resulted in a long lasting reduction in voluntary EtOH consumption in adulthood (Gilpin et al., 2012). It should also be considered that EtOH administrator contact can also be a stressor when evaluating self-administration, making animal-handler habituation important. In addition, placing an animal in an unfamiliar apparatus to perform a task could induce stress or reveal novelty seeking behavior, resulting in higher initial EtOH consumption. In addition, the use of sucrose sweetened EtOH has been shown to confound the assessment of later EtOH consumption (Broadwater et al., 2013). Broadwater et al (2013) showed that voluntary access to 10% EtOH in ‘supersac’ (0.125% saccharin, and 3% sucrose) during adolescence leads to an increase in voluntary consumption of EtOH in ‘supersac’ in adulthood. They also found an increase in supersac consumption in adult rats that consumed only the supersac solution during adolescents. This study and others like it (see (Overstreet et al., 1993) and (Kampov-Polevoy et al., 1990)) reveal that this trend may be the result of solution familiarity or be driven by the reward induced by the sweetened solutions, suggesting that involuntary approaches can confound our insight into complex behaviors that involve subsequent EtOH self-administration. It is becoming ever apparent that we as scientists need to continually expand and refine our experimental tools, so that we are well equipped to make real-world inferences based on our experimental laboratory models.

In an effort to develop a voluntary EtOH delivery system, de Guglielmo and colleagues (2017) adapted a vapor chamber model that would allow rodents to voluntary self-administer EtOH vapor. The apparatus provided voluntary self-administration of EtOH vapor or clean air after the animal triggered one of two nosepokes. Rats underwent 8-hour non-escalating (15 mg/L of EtOH vapor for 2 minutes) or escalating (15mg/L of EtOH vapor for 2, 5, and 10 minutes) voluntary EtOH vapor self-administration during their active, dark cycle. This model demonstrates a novel laboratory approach to induce and maintain EtOH dependence in rats without the use of artificial sweeteners, food reinforcement, water restriction, or forced alcohol administration.

Here we modified the voluntary EtOH self-administration chamber regimen created by de Guglielmo and colleagues to observe how social interactions affect moderate EtOH vapor self-administration in adolescent and adult male and female rats. We used male and female Sprague-Dawley rats beginning at postnatal day 30 (PND30) and postnatal day 70 (PND70) for this experiment. These ages were chosen as PND30 is post-pubescent adolescence in rats and coincides with the time in which human adolescents begin to experiment with alcohol while PND70 is the onset of adulthood. Rats underwent 4-hour sessions of EtOH access every other day, for a total of 40 sessions, with or without their cagemate to determine the impact of social interaction on EtOH exposure. We deviated from Guglielmo’s model and used 20 mg/L EtOH administration in a non-escalating paradigm during 4-hour sessions so that multiple groups of animals could be tested in one day (during their awake cycle) without disruption of their circadian rhythm. These data provide insight into the sexually dimorphic nature of voluntary social EtOH vapor administration in a rat model without the use of sucrose fading or food restriction paradigms.

## METHODS

Eight male and eight female PND24 and PND64 Sprague-Dawley rats were received from Hilltop Laboratories (state, USA), double-housed, and maintained in a temperature- and humidity-controlled room with *ad libitum* access to food and water. Animals were handled daily and allowed to acclimatize for 5 days on a reverse 12-hr/12-hr light:dark cycle (lights off at 6:30 am; lights on at 6:30 pm). All procedures were conducted in accordance with the guidelines of the American Association for the Accreditation of Laboratory Animal Care and the National Research Council’s Guide for Care and Use of Laboratory Animals and approved by the Marshall University IACUC.

### Vapor Chamber Apparatus

The apparatus consists of a 36.83 cm x 25.4 cm x 22.86 cm airtight chamber equipped with two nose pokes located on opposite sides of the chamber and corresponding cue lights positioned immediately above each nose poke (de Guglielmo et al., 2017). One nose poke is active, and one is inactive. The active nose poke when initiated releases ethanol vapor (20 mg/L for 2 mins) into the chamber. The inactive nose poke results in no response. The chamber is ventilated with 15 L/min of air at all times. Nosepoke activity was monitored using infrared beam breaks.

### Chronic intermittent ethanol vapor self-administration

Beginning PND30 and PND70 animals were habituated to the procedure room in their home cages (double housed) for 30 minutes prior to self-administration procedure. Animals were weighed and placed in the vapor chamber for 4 hours every other day during their active, dark, cycle. The active nose poke that initiated EtOH vapor release into the chamber was initially randomized but remained the same throughout the remainder of the sessions. All animals received 10 sessions with their same sex cagemate (voluntary social access, SA) followed by 10 sessions in isolation (voluntary isolated access, IA). All animals then underwent 10 days forced abstinence in which they remained in their homecage (double housed). Following forced abstinence, animals underwent another 10 sessions of isolated access (IA) followed by another 10 sessions of cagemate access (SA), i.e., a reversal of the earlier access paradigm (Figure 1). Ethanol preference was determined by a calculating the percent of active nosepokes (ethanol delivery) out of total nosepokes in each session.

**Figure 1.**
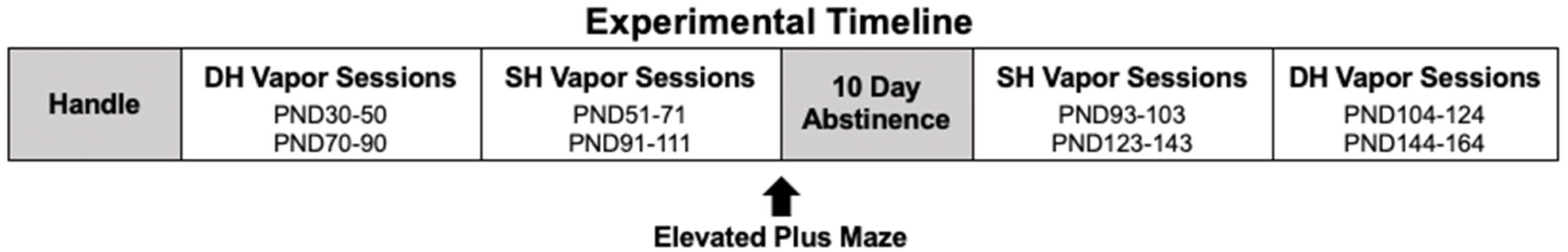
Experimental timeline for voluntary self-administration EtOH vapor chambers. Animals arrived at PND24 and were handled prior to beginning voluntary EtOH self-administration. Beginning on PND30 (adolescents) or PND70 (adults) animals were placed in vapor chambers with their cagemate for 10 sessions voluntary social access (SA). The animals then transitioned to 10 sessions of voluntary isolated access (IA) where they were placed in the chamber without their cagemate. This was followed by a 10-day forced abstinence. Animals then underwent another 10 sessions of IA followed by a final 10 sessions of SA with their cagemate.

### Elevated Plus Maze

To determine changes in anxiety, 24 hr after session 20 animals were tested in the elevated plus maze as previously described in (Wilson et al., 2013). The apparatus consisted of a plus shaped maze made of opaque Plexiglas, two open arms and two closed arms of the same size (50 cm x 10 cm) but with side walls (40 cm). The apparatus was elevated 50.8 cm above the floor. Lighting was set at 35-40 lux, lumen/m2. The rats were habituated to the room for 30 minutes and then placed one at a time in the central area of the maze facing an open arm. Rats were allowed to explore the apparatus for 10 min and then returned to their homecage. Between animals the apparatus was cleaned with 5% vinegar solution. Activity was recorded using an overhead camera. Total number of entries into open versus closed arms and time spent in open versus closed arms was quantified using Anymaze software (Stoelting, IL, USA).

### Withdrawal Score

EtOH behavioral withdrawal signs were assessed 24-hr after the 10th, 20th, 30th, and 40th session (Macey et al., 1996). Abnormal gait, tail rigidity, ventromedial limb retraction, irritability to touch (vocalization), and body tremors were assigned a score of 0-2 based on severity. 0 = no sign, 1 = moderate sign, 2 = severe sign. Signs were summed and used as a quantitative measure of severity of withdrawal.

### Blood Ethanol Concentrations

To avoid possible confounds that may have arisen due to the stress of blood draws and the timing of the animals’ last voluntary EtOH vapor self-administration, a separate group of animals were used for BECs. After completion of the voluntary 40-session self-administration experiment, we assessed the average number of minutes of EtOH vapor per hour per animal per session block was calculated to determine the peak hour of exposure. A separate group of male and female adolescent and adult rats (n=4) were then placed in the vapor chambers for 1 hour. EtOH vapor was passively administered so that the animals received the same amount of EtOH vapor as the peak hour of animals they were representing. Approximately 150 µl of blood was drawn from the lateral saphenous vein at immediately following the EtOH vapor session. Serum was collected from centrifuged samples and stored at −80°C. Samples were analyzed in triplicate using an Analox GL5 alcohol analyzer (Analox Instruments, Lunenburg, MA).

### Statistical Analysis

Comparisons between session, age, and sex were made using repeated measures Analysis of Variance. Post hoc analysis was conducted using Tukey’s multiple comparison test. All analyses were conducted using GraphPad Prism 8 (GraphPad Software, San Diego, California, USA). Statistical significance was assessed using an alpha level of 0.05. All data are presented in figures as the mean +/− S.E.M.

## RESULTS

### % EtOH Preference

Using a non-escalating paradigm to mimic intermittent social drinking these data revealed a significant effect of sex (F_3_, _28_ = 5.5, *p*=0.0042) and session (F_39_, _1092_ = 5.128, *p*<0.0001) across all 40 sessions (Figure 2 A,B). When analyzed based on session block (first SA, first IA, second IA block, and second SA) there was a significant effect of session block (F_3_, _84_ = 20.84, *p*<0.0001) and sex (F_3_, _28_ = 5.5, *p*=0.0042) (Figure 2 C,D). Post hoc analysis revealed that in males exposed to EtOH beginning PND30 there was an increase in EtOH preference (as indicated by an increase in active vs. inactive nosepokes) from the first SA block to the second IA block (*p*=0.0025) and the second SA block (*p*<0.0001). There was also a significant increase from the first IA to the second SA (*p*=0.0039). There was no effect of session block in the female rats exposed to EtOH beginning PND30. In males exposed to EtOH beginning PND70 there was a significant increase in EtOH preference from the first SA block to the second IA block (*p*=0.0002) and second SA block (*p*=0.0003). There was no effect of session block in the female rats exposed to EtOH beginning PND70. These data indicate that there is an increased preference for EtOH across the 40 sessions in male rats regardless of whether exposure to EtOH was initiated in adolescence or adulthood that is not observed in females.

**Figure 2.**
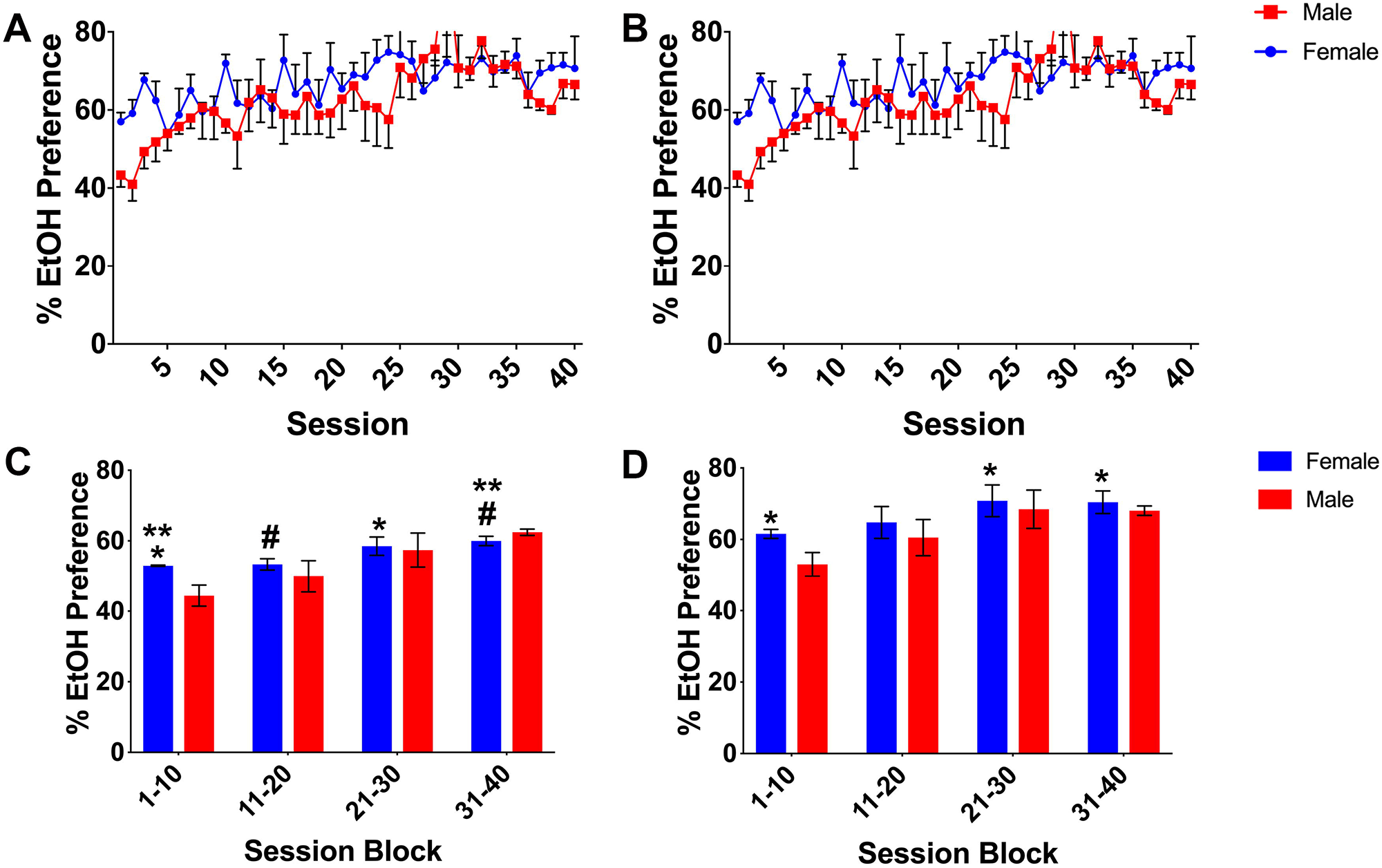
EtOH Preference. Animals underwent 4-hour voluntary self-administration sessions (Figure1) in which they could nosepoke for self-administration of 20 mg/L of EtOH for 2 minutes. %EtOH preference over 40 sessions of voluntary self-administration of EtOH vapor (A) PND30 female and male rats, (B) PND70 female and male rats, (C) summary of EtOH preference per session block in PND30 female and male rats. **\*\*** *p* < 0.01 when comparing PND30 male session block 1-10 to session blocks 21-30 and 31-40. **##** *p* < 0.01 when comparing PND30 male session blocks 11-20 and 31-40. %EtOH preference over 40 sessions of voluntary self-administration of EtOH vapor (D) summary of EtOH preference per session blocks in PND70 female and male rats. ** *p* <0.01 when comparing PND70 male session blocks 1-10, 21-30, and 31-40. ## *p*<0.01 when comparing PND70 male session blocks 1-10 to 31-40. There was a significant overall effect of sex (*p* = 0.0042) and session (*p* < 0.0001) across all 40 sessions that is not indicated in figure. Data is represented as Mean ± SEM, n=8, *=p<0.05.

### Minutes of EtOH Vapor per Body Weight

3-way ANOVA (sex x age x session) revealed a significant effect of sex (F_1, 1120_ = 698.9, p<0.0001), session (F_39, 1120_ =45.88, p<0.0001), and age (F_1, 1120_ = 151.9, p<0.0001) across all 40 sessions (Figure 3 A,B). There was a significant interaction between session x age (F_39, 1120_ =16.88, p<0.0001), session x sex (F_39, 1120_ =1.713, p=0.0045), as well as age x sex (F_1, 1120_ = 4.929, p=0.0266). To further assess the impact of SA versus IA sessions we conducted within group 3-way ANOVA (sex x age x session block). There was an overall effect of sex (F_1, 112_ = 143.9, p<0.0001), session block (F_3, 112_ =84.99, p<0.0001), and age (F_1, 112_ = 31.28, p<0.0001). While there was no significant interaction between sex and age (F_1, 112_ =1.015, p=0.3159), there was a significant interaction between session block x age (F_3, 112_ =26.90, p<0.0001) with a trend towards a session block x sex interaction (F_3, 112_ =3.189, p=0.0713).

**Figure 3.**
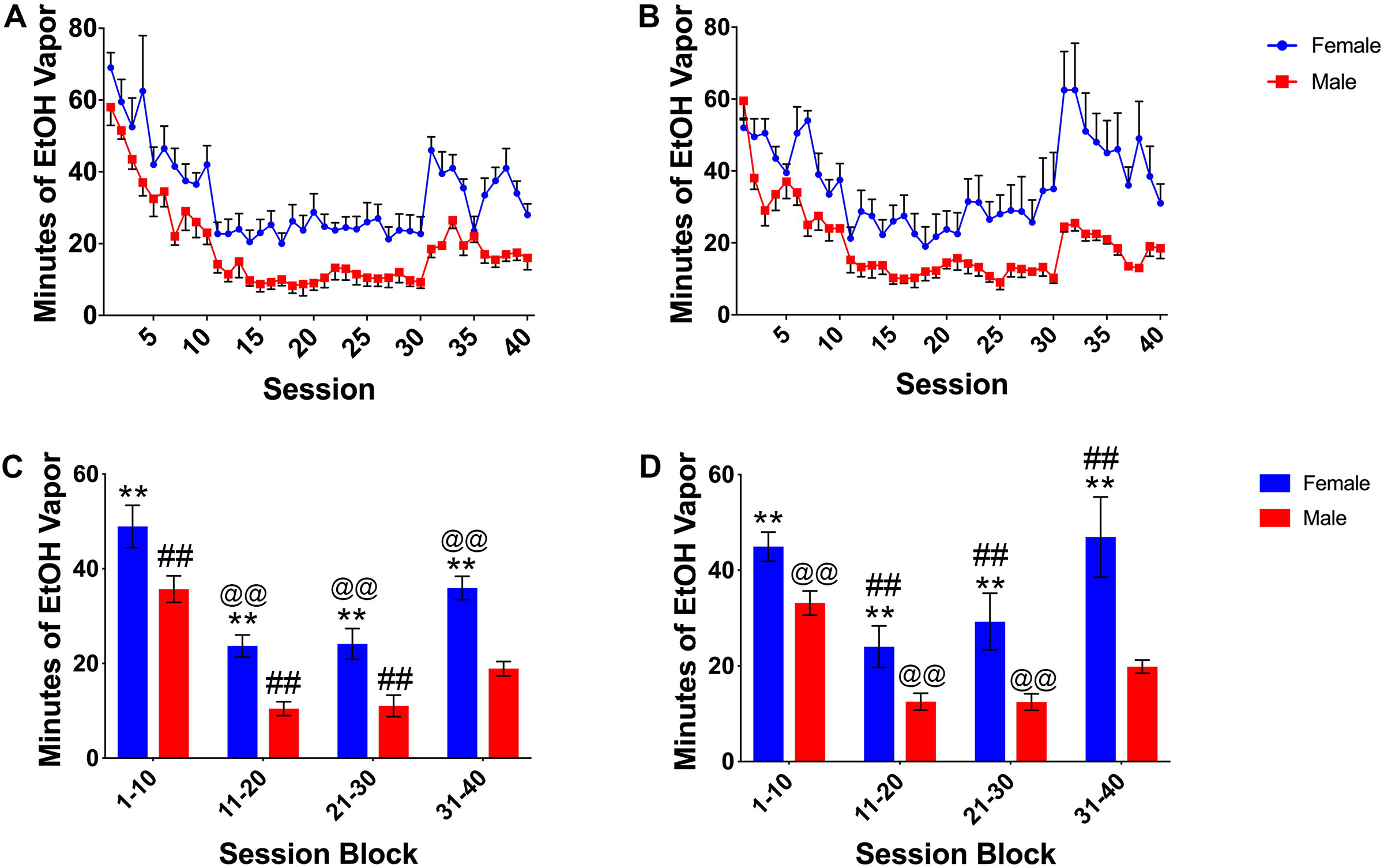
Minutes of EtOH Vapor per Body Weight. Animals underwent 4-hour voluntary self-administration sessions (Figure 1) in which they could nosepoke for self-administration of 20 mg/L of EtOH for 2 minutes. Minutes of EtOH adjusted for body weight over 40 sessions of voluntary self-administration (A) PND30 female and male rats, (B) PND70 female and male rats, (C) minutes of EtOH vapor per body weight summarized across session blocks in PND30 female and male rats. **\*\*** *p* < 0.01 when comparing PND30 female session block 1-10 to the subsequent session blocks 11-20, 21-30, and 31-40. **##** *p* < 0.01 when comparing PND30 male session blocks 1-10, 11-20, 21-30, and 31-40. Minutes of EtOH adjusted for body weight over 40 sessions of voluntary self-administration, (D) minutes of EtOH vapor per body weight summarized across session blocks in PND70 female and male rats. ** *p* <0.01 when comparing PND70 female session block 1-10 to subsequent session blocks 11-20 and 21-30. @ *p* = 0.0456 when comparing PND70 female session blocks 21-30 and 31-40. **##** *p* < 0.01 when comparing PND70 male session blocks 1-10, 11-20, 21-30, and 31-40. There was a significant overall effect of sex (*p* < 0.0001), session (*p* < 0.0001), and age (*p* <0.0001) across all 40 sessions that is not indicated in figure. Data is represented as Mean ± SEM, n=8, *=p<0.05.

Post hoc analysis revealed that female PND30 rats self-administered more EtOH vapor per body weight in their first SA session block when compared to subsequent IA session blocks (first IA block, p<0.0001, second IA block, p<0.0001) as well as the second SA block, p<0.0001, demonstrating a decrease in voluntary EtOH self-administration when female PND30 rats transitioned to the first and second IA session blocks. There was no increase in EtOH self-administration from the first to the second IA session blocks (p=0.8015) despite the intermediate forced abstinence period (Figure 3 C).

Male adolescent rats beginning EtOH vapor exposure at PND30 demonstrated a session block effect. Male rats voluntarily administered more EtOH vapor per body weight in their first SA session block when compared to subsequent session blocks (p<0.0001 (first IA block), p<0.0001 (second IA block), p<0.001 (second SA block)). There was no difference between the first and second IA session blocks (p=0.9777) despite the forced abstinence period (Figure 3 C).

Female PND70 rats voluntarily self-administered more EtOH vapor in their first SA session block when compared to the first IA session block (p<0.0054). There was a significant increase in EtOH exposure when female rats transitioned back to the second SA session when compared to the second IA sessions (p=0.0456). There was no difference between the first and second IA session blocks (p=0.0.7620) despite the intermediate forced abstinence period. There was no significant change EtOH exposure when comparing the first SA session and the second SA session (p=0.6384). Adult males beginning voluntary EtOH vapor exposure at PND70 demonstrated a session block effect. Post hoc analysis within group comparison revealed that the male rats consumed more EtOH in their first SA session block when compared to subsequent session blocks (p=0.0003 (first IA block), p<0.0001 (second IA block), p=0.0006 (second SA block)). There was a significant increase in EtOH vapor self-administration when transitioning back to the second SA block when compared to the second IA block (p=0.0027). There was no difference between the first and second IA session blocks (p=0.8131) despite the forced abstinence period (Figure 3 D).

### Timeout Nose Pokes

To ensure that all animals learned the tasks equally well and to determine whether impulsive nose poking contributes to differences in EtOH consumption we assessed total timeout nose pokes, theorizing that the number of timeout nose pokes should decline as the animals learned the task. 3 way ANOVA (sex x age x session) revealed a significant sex (F_1, 29_ = 26.27, *p*<0.0001), age (F*39, 1092* = 23.78, *p*<0.001), and session effect (F_1,28_ = 11.21, *p*=0.0023) (Figure 4). Post hoc analysis revealed that these effects were mainly driven by differences in timeout nose pokes during early sessions when animals were learning the task. Secondary analysis revealed that female PND30 rats nose poked during timeout significantly more than male PND70 and female PND70 rats during sessions 1-4 (Session 1 p<0.0001, p<0.0001, session 2 p<0.0001, p<0.0001, session 3 p=0.0001, p<0.0001, and session 4 p<0.0001, p<0.0001, respectively) (Figure 4 A). By session 5 there was no difference between groups based on age or sex when compared across identical sessions (p>0.05). As with female PND30 rats, male PND30 rats nosed poked significantly more than PND70 male and female rats during session 1 (p<0.0001, p<0.0001) and session 2 (p<0.0001, p<0.0001) (Figure 4 A). By session 3 there was no difference between groups based on age or sex when compared across identical sessions (p>0.05) suggesting that impulsive nose poking may influence nose poke self-administration of EtOH during the first SA session block as there wasn’t a recapitulation of excess timeout nose pokes in the second SA session block (Figure 4 B). These data suggest that impulsive nosepoking is likely not an underlying factor influencing long-term differences in EtOH self-administration.

**Figure 4.**
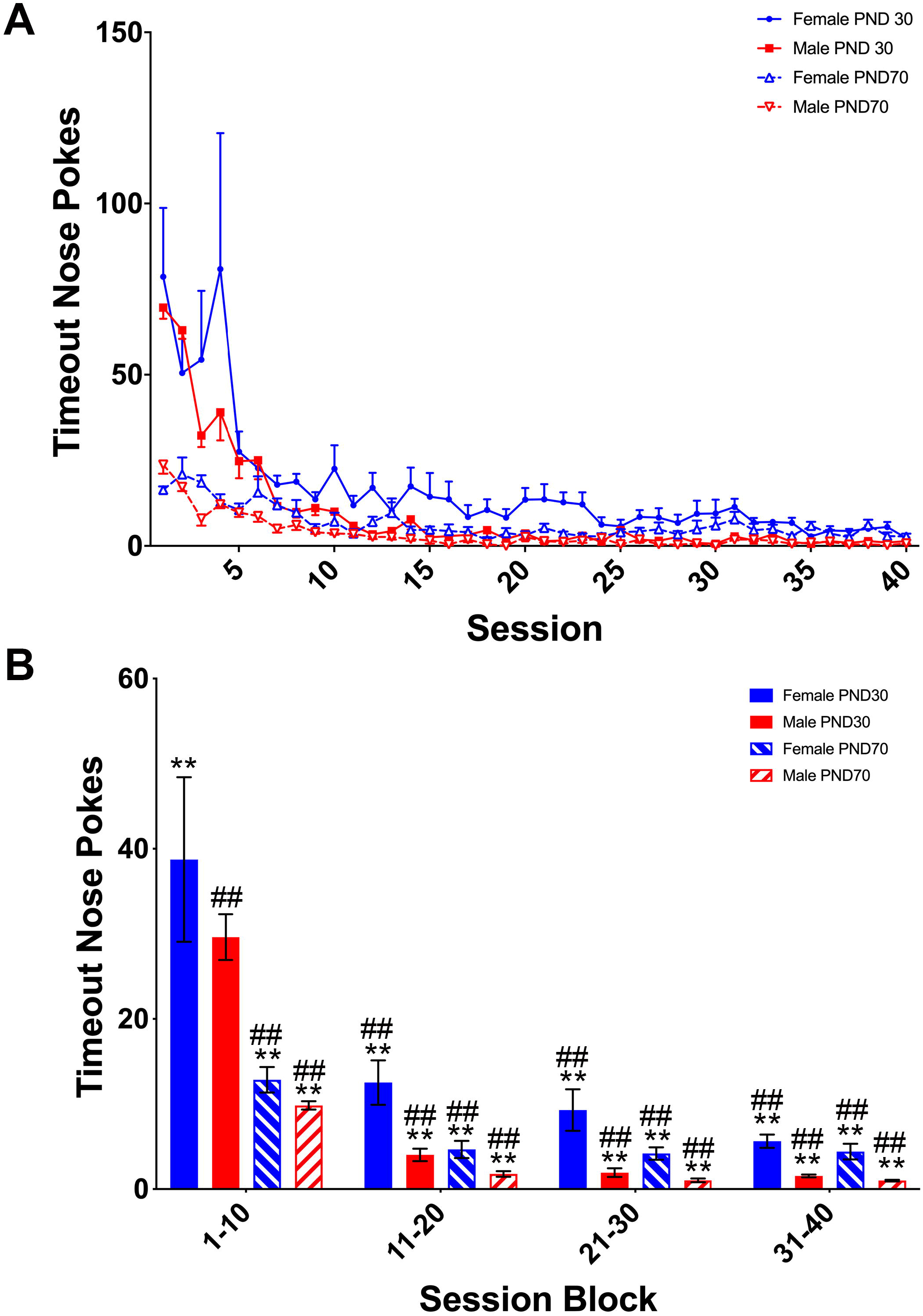
Timeout Nose Pokes. Timeout nosepokes that were initiated during the 2 min of vapor administration were counted across all 40 sessions (A) PND30 and PND70 female and male rats. Impulsive nosepoking decreased in adolescent groups by the 5th session and in adults by the 3rd session. (B) Summary of total nose pokes per session block in PND30 and PND70 female and male rats. ** p<0.01 when comparing PND30 female session block 1-10 to male and female PND70 session block 1-10, as well as PN30 and PND70 male and female session blocks 11-20, 21-30, and 31-40. ## p<0.01 when comparing PND30 male session block 1-10 to male and female PND70 session block 1-10, as well as PN30 and PND70 male and female session blocks 11-20, 21-30, and 31-40. Data is represented as Mean ± SEM, n=8.

### Elevated Plus Maze

To determine whether differences in anxiety may be an underlying factor influencing EtOH consumption we conducted the EPM 24 hours after session 20 (Figure 5). Statistical analysis revealed no significant differences in time spent in open (F_3, 28_=2.878, p=0.0537) and closed arms (F_3, 28_=1.981, p=0.1397) or in distance moved in closed (F_3,28_=1.268, p=0.3043) and open (F_3,28_=2.032, p=0.1321) arms.

**Figure 5.**
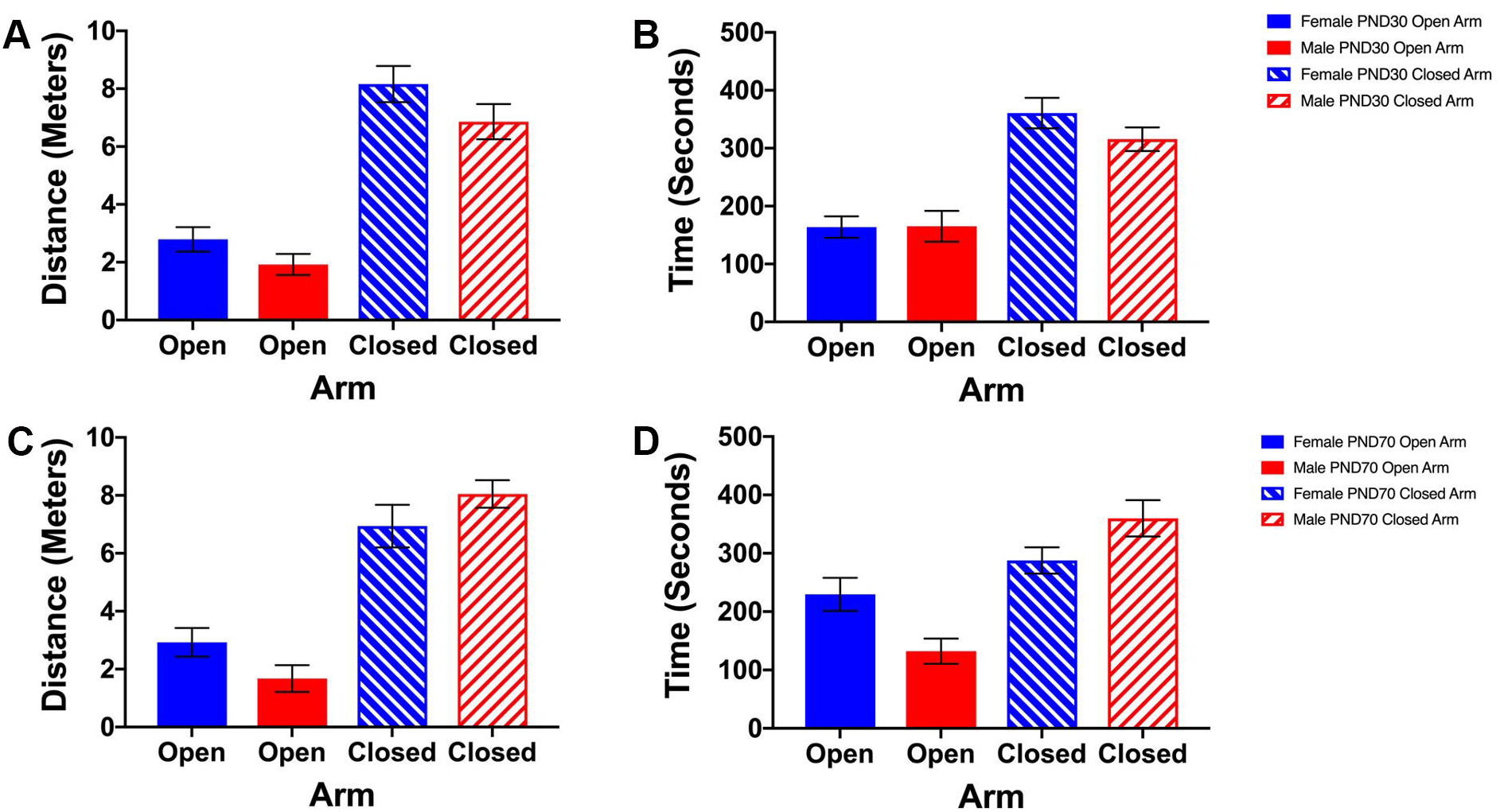
Elevated Plus Maze (EPM). 24hrs after session 20, the period of peak withdrawal, EPM was used to determine changes in anxiety levels. Distance and time spent in open and closed arms of the elevated plus maze. (A) Distance traveled in open and closed arms by PND30 female and male rats. (B) Time spent in open and closed arms by PND30 female and male rats. (C) Distance traveled in open and closed arms by PND70 female and male rats. (D) Time spent in open and closed arms by PND70 female and male rats. Data is represented as Mean ± SEM, n=8, *=p<0.05.

### Somatic Symptoms of Withdrawal

24 hours after the 10^th^, 20^th^, 30^th^, and 40^th^ session rats were scored for withdrawal (Figure 7). All rats showed minimal to moderate signs of withdrawal, however there was no significant effect of sex, age, session, or interaction on overall withdrawal score (p>0.05) (Figure 7 A). When broken down based on separate withdrawal measures there was an overall effect of sex on ventromedial limb retraction (VLR) (F_1,3_=7.658, p=0.0066) however post hoc analysis revealed no individual differences (p>0.05) (Figure 7 B). There was no significant effect of age, sex, or session block in the abnormal gait (AG) (Figure 7 C). There was a significant main effect of age on vocalization (VOC) (F_1,3_=47.91, p<0.0001) (Figure 7 D) and tail rigidity (TR) (F_1,3_=12.1, p=0.0007) (Figure 7 E) and a main effect of session (F_1,3_=5.317, p=0.0018) and age (F_1,3_=4.131, p=0.0445) when body tremors (BT) (Figure 7F) were assessed but post hoc analysis revealed no individual differences (p>0.05).

**Figure 6.**
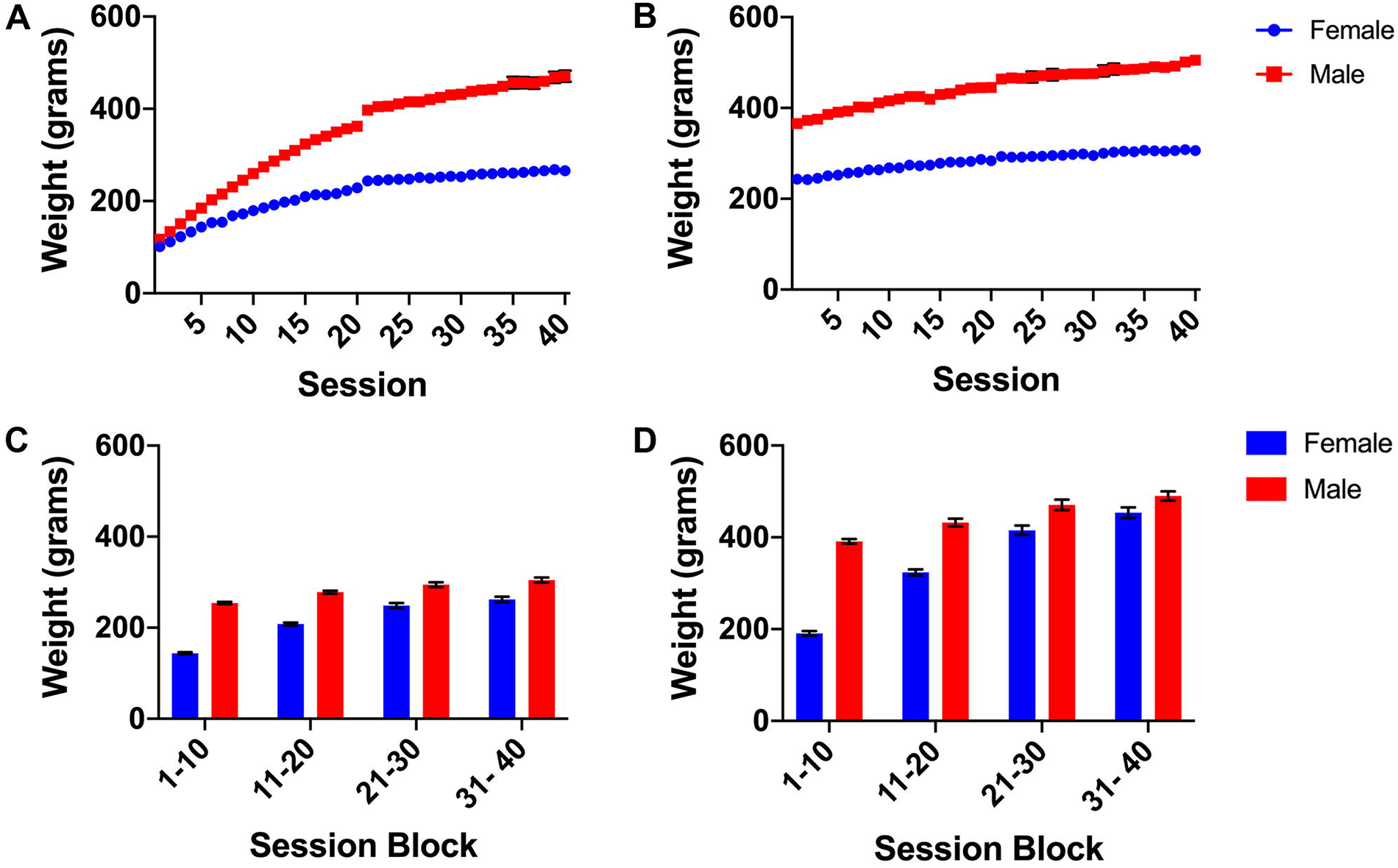
Animal Weights. Summary of PND30 and PND70 female and male rat weights at the beginning of each vapor session. Animal weight progressed naturally showing no sign that the vapor paradigm had a negative impact on growth rate. Data is represented as Mean ± SEM, n=8, *=p<0.05.

**Figure 7.**
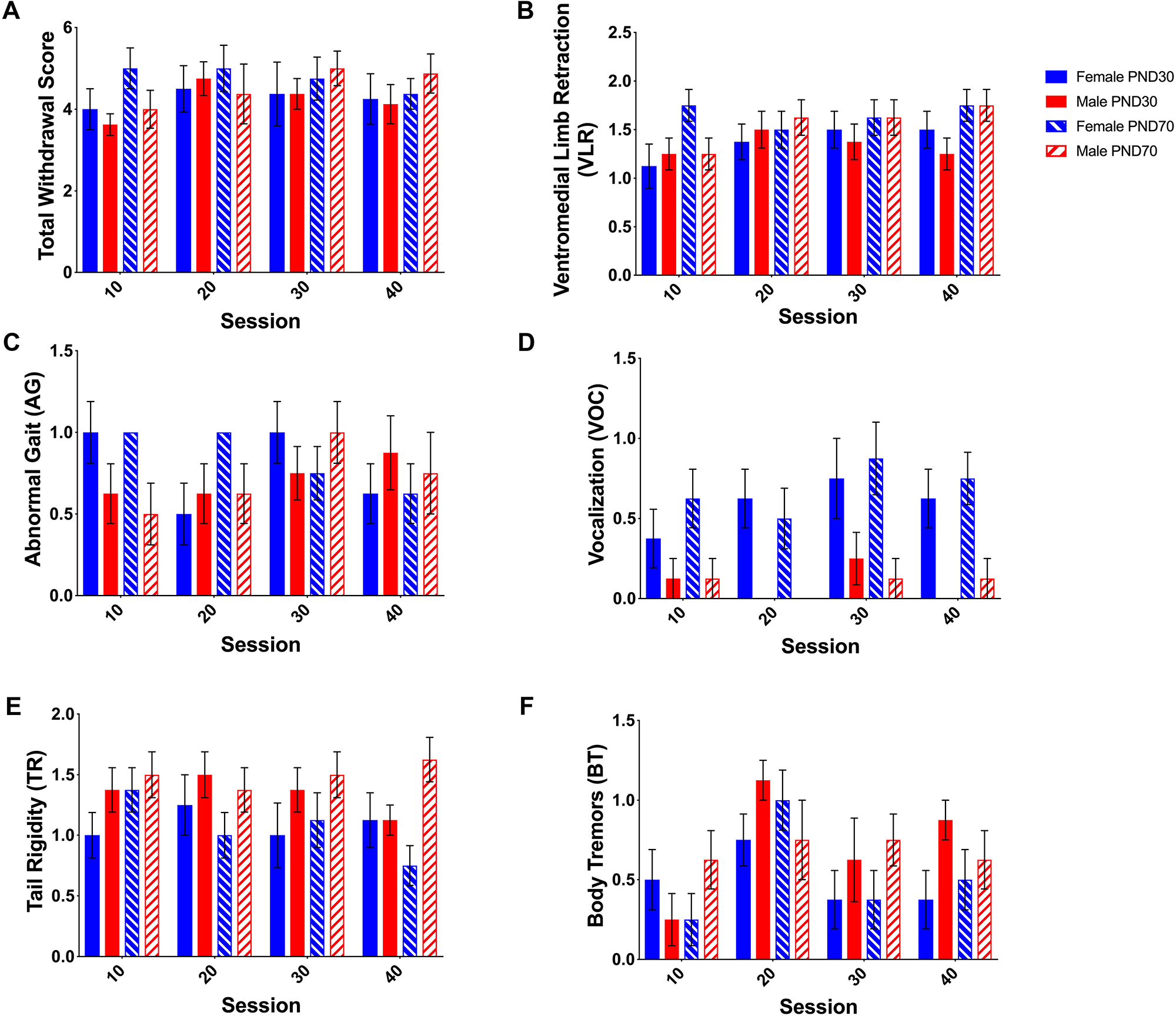
Withdrawal Signs. After each session block, animals were assessed for signs of withdrawal. (A)Total Withdrawal Score, (B) Ventromedial Limb Retraction (VLR), (C) Abnormal Gait (AG), (D) Vocalization (VOC), (E)Tail Rigidity (TR), and (F) Body Tremors (BT) assessment of PND30 and PND70 female and male rats 24 hours after the 10th, 20th, 30th, and 40th voluntary self-administration EtOH vapor sessions. There were no indicators of EtOH withdrawal.

### Weight

There was a significant effect of sex (F_1,39_=5251, p<0.0001), age (F_1,39_=16098, p<0.0001), and session (F_39,39_=215.4, p<0.0001) and a significant interaction between session x sex (F_39,39_=43.16, p<0.0001), session x age (F_39,39_=28.07, p<0.0001), sex x age (F_1,39_=204.8, p<0.0001), and session x sex x age (F_39,39_=6.907, p<0.0001) when assessing weight change (Figure 6). Female and male PND30 rats began sessions weighing the same (p>0.9999) but less than female and male PND70 rats (p<0.001). PND30 male rats continued to grow at a quicker rate than PND70 male rats reaching equivalent weights by session 40 (p=0.8556), whereas female PND30 rat weights remained lower than the female PND70 counterparts (p<0.0001).

### Blood Ethanol Concentrations (BECs)

Due to the voluntary nature of the study, accurate BECs were difficult to determine during the active study as the animals were allowed to voluntarily self-administer EtOH vapor over 4-hour sessions and their drinking patterns across the 4-hour period varied considerably (data not shown). We also wanted to ensure that the additional stress of blood draw did not interfere with the EtOH vapor self-administration. Therefore, a separate group of animals were used for BEC analysis. Animals were placed in the vapor chambers where they received the equivalent average minutes of EtOH vapor as the experimental animals voluntarily administered during their peak hour of exposure (Table 1). Immediately after the 1-hour session, blood was drawn for BEC analysis.

BEC analysis revealed that adolescent (PND30) females achieved an average BEC of 60.02 mg/dl during the first SA session block and 37.26 mg/dl during the second SA session block (Table 1). Adolescent females BECs reached and average of 14.08 mg/dl and 21.72 mg/dl during the first and second IA session blocks, respectively. Male adolescents received lower average minutes of peak EtOH vapor than females across all sessions. Adolescent males achieved average BEC of 44.04 mg/dl in the first SA session. This was not recapitulated in the second SA session as the average BEC was 13.10 mg/dl. Adolescent males achieved average BECs of 5.31 and 10.08 mg/dl in the first IA and second IA.

**Table 1.** Blood EtOH Levels. Average minutes of EtOH for each session block were calculated. Naïve age-matched animals were given passive EtOH to mimic average minutes of EtOH exposure for each session block. BECs were assessed using the analox. Upper panel shows average BECs for adolescent female (left) and male (right). Lower panel shows average BECs for adult female (left) and male (right). Data is represented as Mean ± SEM, n=8, *=p<0.05.

**Adolescent (PND30)**
| **Session Block** | **Females** |  | **Males** |  |
| --- | --- | --- | --- | --- |
|  | **Average Minutes of EtOH** | **BECs (mg/dl)** | **Average Minutes of EtOH** | **BECs (mg/dl)** |
| 1-10 (SA) | 49 | 27.24 | 36 | 33.48 |
| 11-20 (IA) | 24 | 20.35 | 10 | 17.96 |
| 21-30 (IA) | 24 | 20.14 | 10 | 14.63 |
| 31-40 (SA) | 35 | 23.34 | 18 | 18.89 |

**Adult (PND70)**
| **Session Block** | **Females** |  | **Males** |  |
| --- | --- | --- | --- | --- |
|  | **Average Minutes of EtOH** | **BECs (mg/dl)** | **Average Minutes of EtOH** | **BECs (mg/dl)** |
| 1-10 (SA) | 45 | 38.33 | 33 | 36.58 |
| 11-20 (IA) | 24 | 22.39 | 12 | 15.93 |
| 21-30 (IA) | 30 | 24.56 | 12 | 15.69 |
| 31-40 (SA) | 46 | 38.8 | 19 | 16.79 |

Adult (PND70) females achieved an average BEC of 57.34 mg/dl in the first SA session block and 42.44 mg/dl in the second SA session block. The same females reached average BECs of 18.33 and 25.71 mg/dl in the first and second IA sessions, respectively. Adult males reached an average BEC of 35.21 mg/dl during the first SA session. Analysis revealed BECs of 9.27, 13.99, and 17.63 mg/dl after the first IA, second IA, and second SA session blocks.

## DISCUSSION

Over the course of this study male and female adolescent (PND30) and adult (PND70) Sprague-Dawley rats were placed in self-administration vapor chambers for 4-hour sessions on alternate days for a total of 40 sessions. Here we found that males and females, regardless of age do not self-administer high quantities of EtOH vapor. There were no sex by age interactions. However, we found that female rats do voluntarily self-administer more EtOH vapor per body weight than age-matched males and that males increased EtOH preference over session, regardless of age. All animals, regardless of age or sex, voluntarily self-administered more EtOH vapor per body weight during SA sessions than IA sessions however, the BECs remained low throughout, i.e., did not reach binge level ethanol consumption.

Our first goal was to determine if sex and age differences in EtOH preference were consistent with other models of voluntary EtOH self-administration. We did observe an increase in EtOH preference across sessions in males regardless of the age at which EtOH exposure began. This could be because the male rats are still learning the self-administration task, however if this was the case, we would expect to see a similar response in the female rats. This same escalation in EtOH preference was not observed in either female age group. However, female rats at PND30 and PND70 self-administered more EtOH vapor than age-matched males across all session blocks. This is consistent with previous studies demonstrating that females consume more EtOH than males in continuous and intermittent EtOH exposure paradigms using the 2-bottle choice (Priddy et al., 2017, Li et al., 2019). Despite observing a sex effect in EtOH vapor self-administration, we did not see an age-dependent difference in overall EtOH consumed. However, when considering EtOH vapor self-administration relative to body weight we found that adolescent males and females self-administered more EtOH vapor per body weight than their adult counterparts. This is consistent with other studies that have demonstrated that adolescent rats consume more EtOH than adults when adjusted for body weight (Doremus et al., 2005, Hargreaves et al., 2011, Schramm-Sapyta et al., 2014, Vetter-O’Hagen et al., 2009). This is also consistent with studies that have found that there is an increased conditioned taste aversion in adult rats in social settings when compared to adolescents (Vetter-O’Hagen et al., 2009). While we are not orally administering EtOH to these animals, we cannot rule out taste aversion entirely as the animals will be able to taste the EtOH moisture in the chamber and during grooming. However, given that there are no clear pharmacologically relevant BECs this does limit the interpretations based on EtOH exposure

Overall, we found no age-dependent differences in EtOH preference across sessions. Contrary to this finding, Doremus et al. (2005) and Vetter et al. (2007) previously showed that male adolescent rats consume more EtOH than adults in the continuous access 2-bottle choice procedure, however there are three major differences between these studies and the one conducted here. The previous studies did not assess female self-administration so a comparison of females cannot be made. In addition, the BECs obtained from our EtOH vapor studies was much lower (14.63-38.8mg/dL) when compared to that of the previous studies that use oral administration. It is feasible that the differences reported in the previous studies by Doremus et al., (2005) and Vetter et al., (2007) may be due to age-dependent differences in EtOH taste aversion. Schramm-Sapyta et al. (2010) have demonstrated that adolescents display less EtOH taste aversion than adults thus contributing to increased EtOH self-administration. It is possible that we have no age-dependent differences in preference because we are able to bypass this taste aversion when using the EtOH vapor self-administration paradigm and that the age-dependent sensitivity to the rewarding effects of EtOH alone are insufficient to drive an age-effect.

Next, we wanted to determine if social interaction would facilitate changes in a voluntary self-administered EtOH vapor intake. Our model allowed us to observe how social interaction (SA) and isolation (IA) contribute to differences in EtOH voluntary exposure. All rats regardless of age and sex self-administered more EtOH vapor per body weight during the first social session block than subsequent isolated session blocks. While not reaching the same level of minutes of EtOH vapor/body weight that was observed in the first social block, there was a significant increase during the second social block when compared to isolated blocks in the adult females. These results are somewhat consistent with Varlinskaya et al. (2015) who demonstrated a positive correlation between social interaction and consumption of 10% ethanol in “supersac” solution when assessed every other day over six 30-min drinking sessions (3 sessions of social drinking alternating with 3 sessions of drinking alone). During this short access paradigm, male and female adolescent rats and adult male rats all consumed more sweetened EtOH in the social setting versus in isolation. Females displayed a more anxious phenotype, however anxiety levels only correlated with EtOH intake in adolescent females. Unlike our testing paradigm Varlinskaya assessed the correlation between anxiety and social EtOH consumption by placing their rats in an unfamiliar apparatus with unfamiliar partners potentially introducing anxiety as a confound (Varlinskaya and Spear, 2002, Varlinskaya et al., 2015). Higher anxiety has previously been correlated with increase ethanol consumption; however, in this model using cage mates as ‘drinking mates’, we saw no significant age or sex differences when using the EPM, suggesting that anxiety was not a driving factor in EtOH vapor consumption in this task. These data demonstrating that female rats self-administer more EtOH vapor in social environments rather than when alone is consistent with a recent vole study by Anacker et al. (2011). Here, voles were tested for baseline drinking and separated into high and low drinkers. High drinkers were placed with low drinkers, high drinkers with high drinkers, and low drinkers with low drinkers. The authors were able to demonstrate that alcohol consumption was influenced by alcohol consumption of the other animal.

The current data suggests that socialization does not influence EtOH seeking behavior, in this self-administration paradigm, in adolescent male and female rats as indicated by increased self-administration during the first SA session but not the second SA session. The high rate of self-administration of EtOH vapor in the first SA session may be, in part, due to a generalized increase in impulsivity that is observed in adolescent rats when compared to adults (Doremus-Fitzwater et al., 2012). Adolescent rats also tend to have a higher proclivity towards novelty when compared to adult rats (Swartzwelder et al., 2012). Both factors may contribute to the higher EtOH vapor self-administration observed in the first SA session. Reduction in novelty may contribute to the diminished EtOH seeking behavior observed in the second SA session when compared to the first. Another factor to consider would be social play. Social play occurs at higher rates in adolescent rats when compared to adults (Panksepp, 1981) and it is possible that this may also contribute to diminished EtOH seeking behavior in adolescents since they could be spending more time at play rather than consuming EtOH. Additional investigation into how much time rats spend participating in social play in this apparatus would provide further insight into the differences between adolescent and adult EtOH seeking behavior.

Rats are highly social animals suggesting that sociability may be an important factor associated with higher EtOH intake. This is consistent with some human data that demonstrate that a preference for social activity/interaction is associated with higher ethanol intake (Kuntsche et al., 2006, Ham and Hope, 2003). The increased EtOH preference over time in male rats could suggest that sociability may not serve as a strong contributor to the escalation of drinking in both adolescent and adult male rats unlike adult females. Further work will be necessary to determine whether the order of SA and IA sessions contributes to the differences in EtOH self-administration.

## CONCLUSION

It is important to have a diverse set of experimental tools when working in a laboratory so that we may make accurate translational correlations. This novel voluntary self-administration EtOH vapor model developed by La Jolla Alcohol Research Inc. is the first of its kind and allows us to begin to observe the age and sex-dependent effects of socialization in rats in a unique context. Female rats self-administered more EtOH vapor than age-matched males, however male rats regardless of age escalated EtOH voluntary exposure over time. Adolescent and adult rats regardless of sex or age self-administered more EtOH vapor during peer interactions, this was particularly apparent in females. Together, these data provide insight into the sexually dimorphic nature of voluntary EtOH exposure and the impact of social interaction during EtOH access in a rat model. However, one major caveat of this experimental approach is the lack of high BEC’s generated throughout this EtOH vapor self-administration paradigm.

As with any experimental design it is important to select techniques that will work well to garner insight into the hypothesis under investigation. This self-administration EtOH vapor model allows for the investigation into how voluntary self-administration of EtOH vapor changes across adolescence and into adulthood and provides a potential novel method by which to understand the implications of stress and socialization. This method is limited to understanding the equivalent of low-level, casual drinking behavior and not the type of consumption that is associated with escalation, risky behavior, and the development of dependence. This model does allow researchers to observe voluntary EtOH exposure but doesn’t overcome the low-self administration problem that persists with other liquid EtOH self-administration paradigms. However, it does remove confounding factors such as the addition of artificial sweeteners, the stress of oral gavage, or the metabolic stress of diet and water restrictions with the caveat of face validity. As with all new techniques, further examination of EtOH intake and metabolism in this model is required to provide greater guidance for researchers when assessing the suitability of this model for their study.

## ACKNOWLEDGMENTS

We acknowledge the following funding sources: National Institutes of Health (2U54GM104942-3) to MLR and Department of Veterans Affairs Career Development Award (BX002505) to MLR. The funders had no role in study design, data collection and analysis, decision to publish, or preparation of the manuscript. The contents of this manuscript do not represent the views of the National Institutes of Health, Department of Veterans Affairs, or the United States Government.

## REFERENCES

Anacker, A.M., Loftis, J. M. & Ryabinin, A.E. 2011. Alcohol intake in prairie voles is influenced by the drinking level of a peer. Alcohol Clin Exp Res, 35, 1884–90.

Anacker, A.M. & Ryabinin, A.E. 2010. Biological contribution to social influences on alcohol drinking: evidence from animal models. Int J Environ Res Public Health, 7, 473–93.

Anderson, R.I., Varlinskaya, E. I. & Spear, L.P.. 2010. Ethanol-induced conditioned taste aversion in male sprague-dawley rats: impact of age and stress. Alcohol Clin Exp Res, 34, 2106–15.

Arnett, J.J. 2004. Emerging adulthood : the winding road from the late teens through the twenties, New York ; Oxford, Oxford University Press.

Baumrind, D. 1987. A developmental perspective on adolescent risk taking in contemporary America. New Dir Child Dev, 93–125.

Blanchard, R.J., Hori, K., Tom, P. & Blanchard, D.C.. 1987. Social structure and ethanol consumption in the laboratory rat. Pharmacol Biochem Behav, 28, 437–42.

Borsari, B. & Carey, K.B. 2003. Descriptive and injunctive norms in college drinking: a meta-analytic integration. J Stud Alcohol, 64, 331–41.

Broadwater, M., Varlinskaya, E.I. & Spear, L.P. 2013. Effects of voluntary access to sweetened ethanol during adolescence on intake in adulthood. Alcohol Clin Exp Res, 37, 1048–55.

Burnett, S. & Blakemore, S. J. 2009. The development of adolescent social cognition. Ann N Y Acad Sci, 1167, 51–6.

Coleman, L.G., JR., Liu, W., Oguz, I., Styner, M. & Crews, F.T. 2014. Adolescent binge ethanol treatment alters adult brain regional volumes, cortical extracellular matrix protein and behavioral flexibility. Pharmacol Biochem Behav, 116, 142–51.

Collaborators, U.S. B. O. D., Mokdad, A. H., Ballestros, K., Echko, M., Glenn, S., Olsen, H. E., Mullany, E., Lee, A., Khan, A. R., Ahmadi, A., Ferrari, A. J., Kasaeian, A., Werdecker, A., Carter, A., Zipkin, B., Sartorius, B., Serdar, B., Sykes, B. L., Troeger, C., Fitzmaurice, C., Rehm, C. D., Santomauro, D., Kim, D., Colombara, D., Schwebel, D. C., Tsoi, D., Kolte, D., Nsoesie, E., Nichols, E., Oren, E., Charlson, F.J., Patton, G.C., Roth, G. A., Hosgood, H. D., Whiteford, H. A., Kyu, H., Erskine, H. E., Huang, H., Martopullo, I., Singh, J. A., Nachega, J. B., Sanabria, J. R., Abbas, K., Ong, K., Tabb, K., Krohn, K. J., Cornaby, L., Degenhardt, L., Moses, M., Farvid, M., Griswold, M., Criqui, M., Bell, M., Nguyen, M., Wallin, M., Mirarefin, M., Qorbani, M., Younis, M., Fullman, N., Liu, P., Briant, P., Gona, P., Havmoller, R., Leung, R., Kimokoti, R., Bazargan-Hejazi, S., Hay, S. I., Yadgir, S., Biryukov, S., Vollset, S. E., Alam, T., Frank, T., Farid, T., Miller, T., Vos, T., Barnighausen, T., Gebrehiwot, T. T., Yano, Y., Al-Aly, Z., Mehari, A., Handal, A., Kandel, A., Anderson, B., Biroscak, B., Mozaffarian, D., Dorsey, E. R., Ding, E. L., Park, E. K., Wagner, G., Hu, G., Chen, H., Sunshine, J. E., Khubchandani, J., Leasher, J., Leung, J., Salomon, J., Unutzer, J., Cahill, L., Cooper, L., Horino, M., et al. 2018. The State of US Health, 1990-2016: Burden of Diseases, Injuries, and Risk Factors Among US States. JAMA, 319, 1444–1472.

Crews, F.T., Robinson, D. L., Chandler, L. J., Ehlers, C. L., Mulholland, P. J., Pandey, S. C., Rodd, Z. A., Spear, L. P., Swartzwelder, H. S. & Vetreno, R.P.. 2019. Mechanisms of Persistent Neurobiological Changes Following Adolescent Alcohol Exposure: NADIA Consortium Findings. Alcohol Clin Exp Res, 43, 1806–1822.

De Guglielmo, G., Kallupi, M. Cole, MD. & George, O. 2017. Voluntary induction and maintenance of alcohol dependence in rats using alcohol vapor self-administration. Psychopharmacology (Berl), 234, 2009–2018.

Doremus, T.L., Brunell, S. C., Rajendran, P. & Spear, L.P. 2005. Factors influencing elevated ethanol consumption in adolescent relative to adult rats. Alcohol Clin Exp Res, 29, 1796–808.

Doremus-Fitzwater, T.L., Barreto, M. & Spear, L.P.. 2012. Age-related differences in impulsivity among adolescent and adult Sprague-Dawley rats. Behav Neurosci, 126, 735–41.

Galef, B.G., JR., Whiskin, E.E. & Bielavska, E. 1997. Interaction with demonstrator rats changes observer rats’ affective responses to flavors. J Comp Psychol, 111, 393–8.

Gilpin, N.W., Karanikas C. A. & Richardson, H.N.. 2012. Adolescent binge drinking leads to changes in alcohol drinking, anxiety, and amygdalar corticotropin releasing factor cells in adulthood in male rats. PLoS One, 7, e31466.

Grant, B.F., Goldstein, R. B., Saha, T. D., Chou, S. P., Jung, J., Zhang, H., Pickering, R. P., Ruan, W. J., Smith, S. M., Huang, B. & Hasin, D.S.. 2015. Epidemiology of DSM-5 Alcohol Use Disorder: Results From the National Epidemiologic Survey on Alcohol and Related Conditions III. JAMA Psychiatry, 72, 757–66.

Ham, L.S. & Hope, D.A. 2003. College students and problematic drinking: a review of the literature. Clin Psychol Rev, 23, 719–59.

Hargreaves, G.A., Wang, E. Y., Lawrence, A. J. & Mcgregor, I.S.. 2011. Beer promotes high levels of alcohol intake in adolescent and adult alcohol-preferring rats. Alcohol, 45, 485–98.

Jackson, K.M., Sher, K. J. & Park, A. 2005. Drinking among college students. Consumption and consequences. Recent Dev Alcohol, 17, 85–117.

Jones, C.M., Clayton, H. B., Deputy, N. P., Roehler, D. R., Ko, J. Y., Esser, M. B., Brookmeyer, K. A. & Hertz, M.F.. 2020. Prescription Opioid Misuse and Use of Alcohol and Other Substances Among High School Students - Youth Risk Behavior Survey, United States, 2019. MMWR Suppl, 69, 38–46.

Kampov-POLEVOY, A.B., Kasheffskaya, O. P. & Sinclair, J.D. 1990. Initial acceptance of ethanol: gustatory factors and patterns of alcohol drinking. Alcohol, 7, 83–5.

Kuntsche, E., Knibbe, R., Gmel, G. & Engels, R. 2006. Who drinks and why? A review of socio-demographic, personality, and contextual issues behind the drinking motives in young people. Addict Behav, 31, 1844–57.

Li, J., Chen, P., Han, X., Zuo, W., Mei, Q., Bian, E.Y., Umeugo, J. & Ye, J. 2019. Differences between male and female rats in alcohol drinking, negative affects and neuronal activity after acute and prolonged abstinence. Int J Physiol Pathophysiol Pharmacol, 11, 163–176.

Macey, D.J., Schulteis, G., Heinrichs, S. C. & Koob, G.F. 1996. Time-dependent quantifiable withdrawal from ethanol in the rat: effect of method of dependence induction. Alcohol, 13, 163–70.

Montesinos, J., Pascual, M., Pla, A., Maldonado, C., Rodriguez-Arias, M., Minarro, J. & Guerri, C. 2015. TLR4 elimination prevents synaptic and myelin alterations and long-term cognitive dysfunctions in adolescent mice with intermittent ethanol treatment. Brain Behav Immun, 45, 233–44.

Overstreet, D.H., Kampov-Polevoy, A. B., Rezvani, A. H., Murrelle, L., Halikas, J. A. & Janowsky, D.S. 1993. Saccharin intake predicts ethanol intake in genetically heterogeneous rats as well as different rat strains. Alcohol Clin Exp Res, 17, 366–9.

Panksepp, J. 1981. The ontogeny of play in rats. Dev Psychobiol, 14, 327–32.

Peterson, A.C., Silbereisen, R. K. & Sorensen, S. 1996. *Adolescent development: A gloval perspective*, New York, Aldine de Gruyter.

Priddy, B.M., Carmack, S. A., Thomas, L. C., Vendruscolo, J. C., Koob, G. F. & Vendruscolo, L.F. 2017. Sex, strain, and estrous cycle influences on alcohol drinking in rats. Pharmacol Biochem Behav, 152, 61–67.

Ramirez, R.L. & Spear, L.P. 2010. Ontogeny of ethanol-induced motor impairment following acute ethanol: assessment via the negative geotaxis reflex in adolescent and adult rats. Pharmacol Biochem Behav, 95, 242–8.

Risher, M.L., Fleming, R. L., Risher, W. C., Miller, K. M., Klein, R. C., Wills, T., Acheson, S. K., Moore, S. D., Wilson, W. A., Eroglu, C. & Swartzwelder, H.S.. 2015a. Adolescent intermittent alcohol exposure: persistence of structural and functional hippocampal abnormalities into adulthood. Alcohol Clin Exp Res, 39, 989–97.

Risher, M.L., Sexton, H. G., Risher, W. C., Wilson, W. A., Fleming, R. L., Madison, R. D., Moore, S. D., Eroglu, C. & Swartzwelder, H.S.. 2015b. Adolescent Intermittent Alcohol Exposure: Dysregulation of Thrombospondins and Synapse Formation are Associated with Decreased Neuronal Density in the Adult Hippocampus. Alcohol Clin Exp Res, 39, 2403–13.

Sacks, J.J., Gonzales, K. R., Bouchery, E. E., Tomedi, L. E. & Brewer, R.D.. 2015. 2010 National and State Costs of Excessive Alcohol Consumption. Am J Prev Med, 49, e73–e79.

Schramm-Sapyta, N.L., Difeliceantonio, A. G., Foscue, E., Glowacz, S., Haseeb, N., Wang, N., Zhou, C. & Kuhn, C.M. 2010. Aversive effects of ethanol in adolescent versus adult rats: potential causes and implication for future drinking. Alcohol Clin Exp Res, 34, 2061–9.

Schramm-Sapyta, N.L., Francis, R., Macdonald, A., Keistler, C., O’Neill, L. & Kuhn, C.M.. 2014. Effect of sex on ethanol consumption and conditioned taste aversion in adolescent and adult rats. Psychopharmacology (Berl), 231, 1831–9.

Sebastian, C., Burnett, S. & Blakemore, S.J. 2008. Development of the self-concept during adolescence. Trends Cogn Sci, 12, 441–6.

Silveri, M.M. & Spear, L.P.. 1998. Decreased sensitivity to the hypnotic effects of ethanol early in ontogeny. Alcohol Clin Exp Res, 22, 670–6.

Slade, T., Chapman, C., Swift, W., Keyes, K., Tonks, Z. & Teesson, M. 2016. Birth cohort trends in the global epidemiology of alcohol use and alcohol-related harms in men and women: systematic review and metaregression. BMJ Open, 6, e011827.

Spear, L.P. & Swartzwelder, H.S. 2014. Adolescent alcohol exposure and persistence of adolescent-typical phenotypes into adulthood: a mini-review. Neurosci Biobehav Rev, 45, 1–8.

Stahre, M., Roeber, J., Kanny, D., Brewer, R.D. & Zhang, X. 2014. Contribution of excessive alcohol consumption to deaths and years of potential life lost in the United States. Prev Chronic Dis, 11, E109.

Swartzwelder, N.A., Risher, M. L., Abdelwahab, S. H., D’abo, A., Rezvani, H., Levin, E. D., Wilson, W. A., Swartzwelder, H. S. & Acheson, S.K.. 2012. Effects of ethanol, Delta(9)-tetrahydrocannabinol, or their combination on object recognition memory and object preference in adolescent and adult male rats. Neurosci Lett, 527, 11–5.

Varlinskaya, E.I. & Spear, L.P. 2002. Acute effects of ethanol on social behavior of adolescent and adult rats: role of familiarity of the test situation. Alcohol Clin Exp Res, 26, 1502–11.

Varlinskaya, E.I., Truxell, E. M. & Spear, L.P. 2015. Ethanol intake under social circumstances or alone in sprague-dawley rats: impact of age, sex, social activity, and social anxiety-like behavior. Alcohol Clin Exp Res, 39, 117–25.

Vetter, C.S., Doremus-Fitzwater, T. L. & Spear, L.P.. 2007. Time course of elevated ethanol intake in adolescent relative to adult rats under continuous, voluntary-access conditions. Alcohol Clin Exp Res, 31, 1159–68.

Vetter-O’Hagen, C., Varlinskaya, E. & Spear, L. 2009. Sex differences in ethanol intake and sensitivity to aversive effects during adolescence and adulthood. Alcohol Alcohol, 44, 547–54.

White, A., Castle, I.J., Chen, C. M., Shirley, M., Roach, D. & Hingson, R. 2015. Converging Patterns of Alcohol Use and Related Outcomes Among Females and Males in the United States, 2002 to 2012. Alcohol Clin Exp Res, 39, 1712–26.

Wilsnack, R.W., Wilsnack, S. C., Kristjanson, A. F., Vogeltanz-Holm, N. D. & Gmel, G. 2009. Gender and alcohol consumption: patterns from the multinational GENACIS project. Addiction, 104, 1487–500.

Wilson, C.A., Vazdarjanova, A. & Terry, A. V., JR. 2013. Exposure to variable prenatal stress in rats: effects on anxiety-related behaviors, innate and contextual fear, and fear extinction. Behav Brain Res, 238, 279–88.

Witt, E. D. 2007. Puberty, hormones, and sex differences in alcohol abuse and dependence. Neurotoxicol Teratol, 29, 81–95.

Wolffgramm, J. & Heyne, A. 1991. Social behavior, dominance, and social deprivation of rats determine drug choice. Pharmacol Biochem Behav, 38, 389–99.

